# CellVGAE: An unsupervised scRNA-seq analysis workflow with graph attention networks

**DOI:** 10.1101/2020.12.20.423645

**Authors:** David Buterez, Ioana Bica, Ifrah Tariq, Helena Andrés-Terré, Pietro Liò

## Abstract

Currently, single-cell RNA sequencing (scRNA-seq) allows high-resolution views of individual cells, for libraries of up to (tens of) thousands of samples. In this study, we introduce the use of graph neural networks (GNN) in the unsupervised study of scRNA-seq data, namely for dimensionality reduction and clustering. Motivated by the success of non-neural graph-based techniques in bioinformatics, as well as the now common feedforward neural networks being applied to scRNA-seq measurements, we develop an architecture based on a variational graph autoencoder with graph attention layers that works directly on the connectivity of cells. With the help of three case studies, we show that our model, named CellVGAE, can be effectively used for exploratory analysis, even on challenging datasets, by extracting meaningful features from the data and providing the means to visualise and interpret different aspects of the model. Furthermore, we evaluate the dimensionality reduction and clustering performance on 9 well-annotated datasets, where we compare with leading neural and non-neural techniques. CellVGAE outperforms competing methods in all 9 scenarios. Finally, we show that CellVGAE is more interpretable than existing architectures by analysing the graph attention coefficients. The software and code to generate all the figures are available at https://github.com/davidbuterez/CellVGAE.

## 1 Introduction and motivation

scRNA-seq allows gene expression to be quantified at the level of individual cells, and the ability to carry out these measurements in a high-throughput fashion was in itself a revolutionary step forward. Single-cell sequencing has its own particular challenges, as technical and biological limitations contribute to noisier and more complex data than previous sequencing techniques [CNS19]. Additionally, different sequencing protocols come with unique advantages and disadvantages, and often to bespoke bioinformatics tools for each protocol. Nevertheless, the potential of scRNA-seq has prompted the development of computational analysis techniques since the inception of the technique in 2009 [TBW^+^09], with the sheer dimensionality and nonlinearity of the data proving challenging even today. Tasks usually tackled by computational scientists are quality control, normalisation, dimensionality reduction, feature selection, cell clustering, trajectory inference, differential expression, gene regulatory network reconstruction, gene dynamics and gene set analysis [CNS19]. A recent review [LT19] identifies 385 scRNA-seq analysis tools as of 7 March 2019, reflecting rapid iterative progress in the field. We observe that non-neural tools are the most widespread, most likely for their simplicity and interpretability; we offer a short overview of relevant techniques in Section 2.2. In general, non-neural methods employ simple but powerful handcrafted algorithms, striking a balance between interpretability and predictive power. As such, we recognise the need not only for optimal task performance, but also for transparency and explainability.

In this work, we investigate a machine learning approach with applications to dimensionality reduction and clustering. Based on the recent interest in both graph-based scRNA-seq clustering and graph neural networks (Sections 2 and 3), we propose a neural model that is built upon the graph variational autoencoder (VGAE) with graph attention layers (GAT), named CellVGAE. Compared to other neural models which work directly on the gene expression values, CellVGAE leverages the connectivity between cells, represented as a graph, to perform convolutions on a non-Euclidean structure, thus subscribing to the geometric deep learning paradigm. We use *k*-nearest neighbour (KNN) and Pearson correlation graphs, based on their widespread use and efficient implementations (especially KNN, e.g. [CFS09], [HAYSZ11], [FC16]).

There are multiple reasons motivating the introduction of a graph neural network methodology. Firstly, the graph can support learning, acting as a valuable inductive bias, by allowing the model to exploit relationships that are impossible or harder to model by the simpler dense layers. This idea has been studied in a different setting by Dutil et al. [DCW^+^18], where protein-protein interaction networks (existing biological knowledge) are used as the graph together with RNA-seq gene expression measurements. On the same note, the message passing paradigm of GNNs allows neighbouring cells to directly exchange and aggregate information in *k*-hops (*k* graph convolution layers). Secondly, graphs are generally more interpretable and visualisable; the GAT framework made important steps in bringing these desirable features to machine learning, a trait not shared by traditional (non-graph) methods. We dedicate the entirety of Section 5.4 to the study of these properties. Thirdly, by jointly using the variational autoencoder and graph neural networks, we allow future studies to exploit advances in both of these areas, which are active research subjects (see Section 6 for an example).

We validate the proposed novel workflow with the help of three case studies, firstly by analysing the clustering performance on a recent scRNA-seq dataset of *Schistosoma mansoni*, proven difficult to analyse by existing methods due to its subtle signals and where CellVGAE can successfully recover the clustering structure and marker genes (Section 5.1). The second case study examines a similarly difficult macrophages dataset in Appendix A. Thirdly, we demonstrate that our method is consistent in identifying quality clusters by applying it to the well-known PBMC3k dataset (Section 5.2). We follow with an extensive analysis where we compare with state-of-the-art methods on multiple clustering metrics, including adjusted rand index and silhouette coefficient, and show leading performance (Section 5.3). Finally, we offer multiple ways of interpreting the learnt properties by leveraging the parameters of the neural network, including some that are unique to CellVGAE, in the form of graph attention coefficients (Section 5.4).

## 2 Related work

Most scRNA-seq analysis workflows require dimensionality reduction for denoising and visualisation of data, for which they often use traditional machine learning techniques such as principal component analysis (PCA); however the standard choice are non-linear algorithms such as t-distributed stochastic neighbour embedding (t-SNE) [vdMH08] and uniform approximation and projection (UMAP) [MHM20]. t-SNE imposes a trade-off between representing global and local features by favouring local structure. Using t-SNE effectively is also complicated by the choice of its *perplexity* parameter, which can significantly alter its behaviour, for example in the number and shape of clusters. UMAP, in comparison, offers a simpler interface and is known to be scalable to a large number of cells. It is recommended as the default option for scRNA-seq studies [LT19]. While the ease-of-use is attractive, these methods are not sufficient on their own. t-SNE, for example, is not recommended for general dimensionality reduction tasks (i.e. other tasks than visualisation in 2 or 3 dimensions) due to uncertainties in the ability to represent the local structure in high dimensional spaces. Additionally, some assumptions of t-SNE, like the local linearity on the manifold can be violated for certain datasets. Thus, the recommendation is to apply t-SNE for visualisation following other dimensionality reduction techniques, such as autoencoders, which are more suited to learning on highly-varying manifolds than local methods [vdMH08]. UMAP also comes with concerns about the preference for local structure and susceptibility to noise [MHM20]. Furthermore, there is empirical evidence confirming that methods such as neural networks outperform t-SNE and UMAP used solely for dimensionality reduction on scRNA-seq data [BATCL20], [LMK20].

### 2.1 Neural network models

Deep learning could be considered a natural step forward in the analysis of biological, and hence, scRNA-seq data, based on its success in fields like computer vision [VDDP18], natural language processing [TSK^+^20], as well as others [Sej20]. However, due to challenging data characteristics (sparsity) and generally low interpretability of deep learning, neural dimensionality reduction approaches are less standard despite their theoretical performance. One of the first studies that implemented this idea is [LJKBJ17]. A significant difference to ours is that the training procedure is largely supervised, with an initial pre-training step done using denoising autoencoders [VLBM08]. Another significant aspect of the architecture is the use of fully-connected dense layers in conjunction with layers that connect only a subset of the neurons, by using known protein-protein interactions and protein-DNA interactions from [SDG^+^12] to connect only certain gene inputs with the hidden neurons (another case of biological inductive bias). The architecture showed competitive performance in inferring cell type for its time. More recently, most of the efforts have been directed at autoencoders (AE) and their variational counterpart (VAE). Out of the existing works, we mention AutoImpute [TMSM18], scVAE [GVT^+^20], Deep Count Autoencoder [ESM^+^19], scIV [SGYP20] and DiffVAE [BATCL20] noting that each of the methods focuses on different challenges.

Out of the neural methods, DiffVAE is the most recent and shows state-of-the-art performance on tasks of dimensionality reduction and clustering. Its motivation is the study of disentanglement and perturbation in the context of cell differentiation. At the core of the method is a Maximum Mean Discrepancy Variational Autoencoder (MMD-VAE), a member of the InfoVAE family of autoencoders introduced in [ZSE17]. The MMD loss fixes two problems of the usual VAE (based on the Evidence Lower Bound): overestimation of latent variance and uninformative latent information. In DiffVAE, the MMD-VAE is combined with t-SNE, *k*-means or DBSCAN [HPD19] for visualisation and clustering. DiffVAE reaches state-of-the-art performance on clustering tasks, evaluated using the Adjusted Rand Index (ARI).

DiffVAE also innovates by showing that neural models can be interpretable. For each of the latent dimensions, the cells that are at least a standard deviation from the mean are selected, and the participation of the identified cells in each of the clusters found at the previous step is computed as a percentage. The top 10 percentages are used to select the most relevant latent dimensions for each cluster. This is combined with layer weights analysis: the weight matrices of the VAE decoder are multiplied, thus enabling a connection between genes and latent dimensions. The genes corresponding to the highest weights for all clusters can thus be interpreted as “marker genes”, with biomedical literature supporting the findings.

### 2.2 Graph-based models

Graph-based clustering on scRNA-seq data was popularised by Seurat [SBH^+^19]. Typically, this involves a *k*-nearest neighbour search combined with a community detection algorithm, such as Louvain clustering [BGLL08]. Efficient implementations exist for both techniques, thus one advantage of graph-based clustering is its scalability. Other positives are that graph construction and clustering operations do not assume that the cells in clusters are normally distributed (as in Gaussian mixture models) and do not favour spherical clusters (*k*-means) [ALB^+^20]. Seurat uses either KNN (*k*-nearest neighbour) or SNN (shared nearest neighbour) graphs.

We have noticed that algorithms that operate on the connectivity of cells (i.e. graphs) are increasingly common. In [ZZX^+^20], Zhu et al. recently introduced the structural shared nearest neighbour-Louvain (SSNN-Louvain) method, which improves upon the simpler KNN or SNN by integrating the structural information of the graph and community module detection. Improvements of the method include reduced parameter tuning and unsupervised detection of clusters. Mohammadi et al. very recently introduced ACTIONet [MDVK20], a computational framework based on multiresolution decomposition to learn dominant transcriptional patterns and characterise cell types and states. Internally, ACTIONet uses a density-dependent nearest-neighbour algorithm (*k^∗^*-NN) to connect cells based on the square root of Jensen-Shannon divergence as metric, but also considering the density of the cell’s neighbourhood. This density-based procedure avoid the specification of the *k* number of neighbours.

Finally, Tarashansky et al. introduced self-assembling manifolds (SAM) [TXL^+^19], an iterative graph-based approach to feature selection. The SAM algorithm starts with a random KNN graph and averages the expression of each cell with its *k* neighbours by performing a weighted 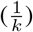 product of the graph adjacency matrix and gene expression matrix. Using the averaged expression, SAM computes a spatial dispersion factor for each gene (gene weights), which are in turn used to rescale the expression matrix. Pairwise distances between cells are recomputed and the *k*-nearest neighbours are updated; the whole procedure repeats until the gene weights converge. The main contribution of SAM is the iterative feature selection procedure, that can theoretically overcome the standard highly variable genes selection (variable genes might not be biologically interesting) coupled with the KNN-based graph generation.

In terms of graph neural network approaches, a recent publication by Ravindra et al. [RSP^+^20] used graph attention networks on scRNA-seq data in a supervised way for predicting disease state of multiple sclerosis, with leading performance compared to other machine learning methods. On a different note, DiffVAE briefly mentions the potential of the graph variational autoencoder for predicting edges on a graph built by Pearson correlation (each cell is connected to the single most similar cell), suggesting possible cell fates.

With a large number of techniques published in a relatively short time, we formulate two criteria for selecting the algorithms that we include in our evaluation of CellVGAE: (1) achieving state-of-the-art performance on dimensionality reduction and clustering (unsupervised learning), while preferably comparing against a large number of methods for a richer and more meaningful evaluation; (2) open source availability of the code and ease of use (ensured by quality documentation) – this is especially important as we include additional metrics in our evaluation, apart from ARI, such as silhouette coefficient and number of found clusters, meaning that we need to be able to benchmark the methods ourselves. Out of the recent publications, we select SAM and DiffVAE as meeting the above criteria.

## 3 Preliminaries

In this work, we assume simple graphs (undirected, unweighted, without loops or multiple edges), defined as a tuple 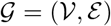, where 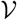 is the set of vertices or nodes {*v*_0_, *v*_1_,…} and 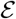 is the set of edges between nodes, 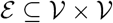. A common representation is given by a graph’s adjacency matrix **A** (whose elements *a_ij_* = 1 if 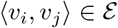, with *a_ij_* = 0 otherwise). For graph neural networks, we also assume that *D*-dimensional node features are represented by a *N* × *D* matrix **X**, where 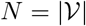.

### Graph variational autoencoder

The graph variational autoencoder (VGAE) is an unsupervised framework introduced by Kipf and Welling in [KW16]. Like the standard VAE, the VGAE has two components: an encoder and a decoder, which are trained to learn latent variables **z**, aggregated in an *N* × *L* matrix **Z**, where *L* is the number of latent dimensions. The encoder or inference model is similar to the VAE, but does not depend only on the signal **X**, but also the connectivity of the graph **A**:

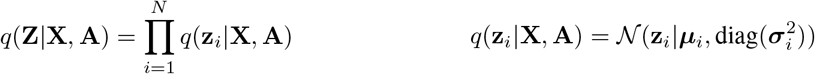

In the original formulation, the parameters ***μ***, ***σ*** are learnt by graph convolutional networks (GCN); formally ***μ*** = GCN_*μ*_(**X**, **A**) and log ***σ*** = GCN_*σ*_(**X**, **A**), where **X** are features learned by previous convolutional layers.

The decoder, or generative model, simply reconstructs an adjacency matrix using the inner product of latent variables:

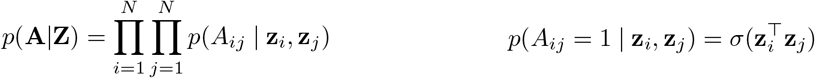

Here, *σ* is the logistic sigmoid function. Also note that only the graph structure is reconstructed (not the node features). The loss function is of the form:

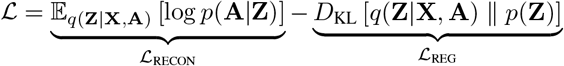

where the two components are the reconstruction loss 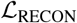 and a regularisation term 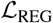. In the standard VGAE framework, the regularisation is given by the Kullback-Leibler divergence between *q*(·) and *p*(·).

### Graph attention networks

Graph attention networks (GAT) [VCC^+^18] are a powerful neural framework for graph-structured data, initially designed for supervised and semi-supervised learning. The main contribution is the introduction of the *graph attention layer*, capable of performing self-attention on graph nodes. Formally, given a set of node features **h** = {*h*_1_, *h*_2_,…, *h_N_*} with *h_i_* ∈ ℝ^*D*^, the first step is to apply a shared linear transformation, parameterised by a weight matrix **W**, on all nodes. This is followed by an attention mechanism *a* applied pairwise to the scaled node features: *e_ij_* = *a*(**W***h_i_*, **W***h_j_*), where **W** ∈ ℝ^*D*′×*D*^ and *a*: ℝ^*D*′^ × ℝ^*D*′^ → ℝ assuming *D*′ is the output dimension of the nodes. The number *e_ij_* can be interpreted as the contribution of node *j*’s features to node *i*. In order to make use of the graph structure, the above computations are limited to the neighbourhood 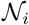 for some node *i*. Finally, the outputs are normalised using the softmax function, resulting in the final form of the attention coefficients:

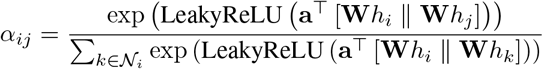

where ^⊤^ is transposition and ‖ is concatenation. The normalised coefficients are used together with a learnable linear combination of the neighbouring features (usually after applying a non-linearity *σ*) to learn the output node features. This can be extended to applying the attention mechanism independently *K* times, a procedure named *multi-head attention*, in two ways:

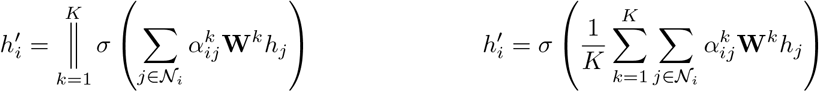

The formulation on the left is appropriate for the hidden (inner) layers as it outputs the concatenated hidden representations of dimension *K* · *D*′. On the other hand, the calculation on the right performs the mean of all the attention heads and is appropriate for the last (output) layer of the network.

## 4 Methods

In this section, we give a complete description of the neural architecture of CellVGAE, followed by an overview of our unsupervised analysis workflow.

### 4.1 CellVGAE architecture

We consider the standard variational graph autoencoder as the starting point. Apart from their demonstrated performance, we choose to use GAT layers instead of GCN for multiple reasons. Firstly, the attention coefficients generated during learning are valuable assets for improving the interpretability of the model, as the authors suggest in the final section of the original study. Since the coefficients indicate the contribution of nodes in a pairwise manner, they can be visualised as a weighted graph between the cells (Section 5.4). Secondly, as GAT layers use a linear transformation under the hood, a traditional layer weights analysis can be carried out, in a manner very similar to DiffVAE, as we show in Section 5.1.2. Finally, graph attention networks have not been studied in unsupervised settings, so we consider this an interesting direction.

We propose an encoder based on graph attention layers. Using the established notation, where **X**_0_ is the initial set of node features and 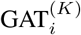 represents the *i*^th^ layer with *K* attention heads, we give the definition of the first and subsequent inner neural layers for an architecture with *N* inner layers:

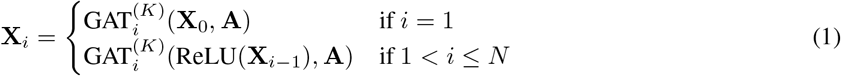

The inner layers concatenate the representations learned by the multi-head attention, so after the first layer the number of output dimensions is *K* · *D_i_* where *D_i_* is the output dimension for each layer. In theory, the number of heads *K* can be different for each layer, but for a simpler architecture we use the same *K* for all layers. The final two layers that learn the parameters ***μ***, ***σ*** follow the second branch of definition (1), i.e.

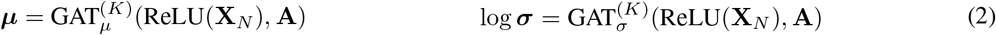

with the exception that they average the features learned by the attention heads instead of concatenating them.

We use the same inner product decoder as the original VGAE, as well as the standard loss that combines regularisation and reconstruction terms:

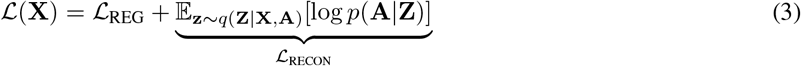

with the mention that 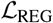 can use any appropriate loss from the literature that minimises the divergence between the learned embeddings and prior distributions. In this work, we experiment with the MMD and KL losses:

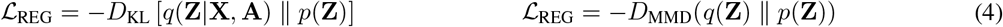

### 4.2 CellVGAE workflow

Having established the neural architecture, we propose an unsupervised scRNA-seq analysis workflow that exploits the strength of CellVGAE for dimensionality reduction and clustering, with the possibility to supplement other downstream analyses (Section 5). We include a simplified overview in Figure 1.

**Figure 1:**
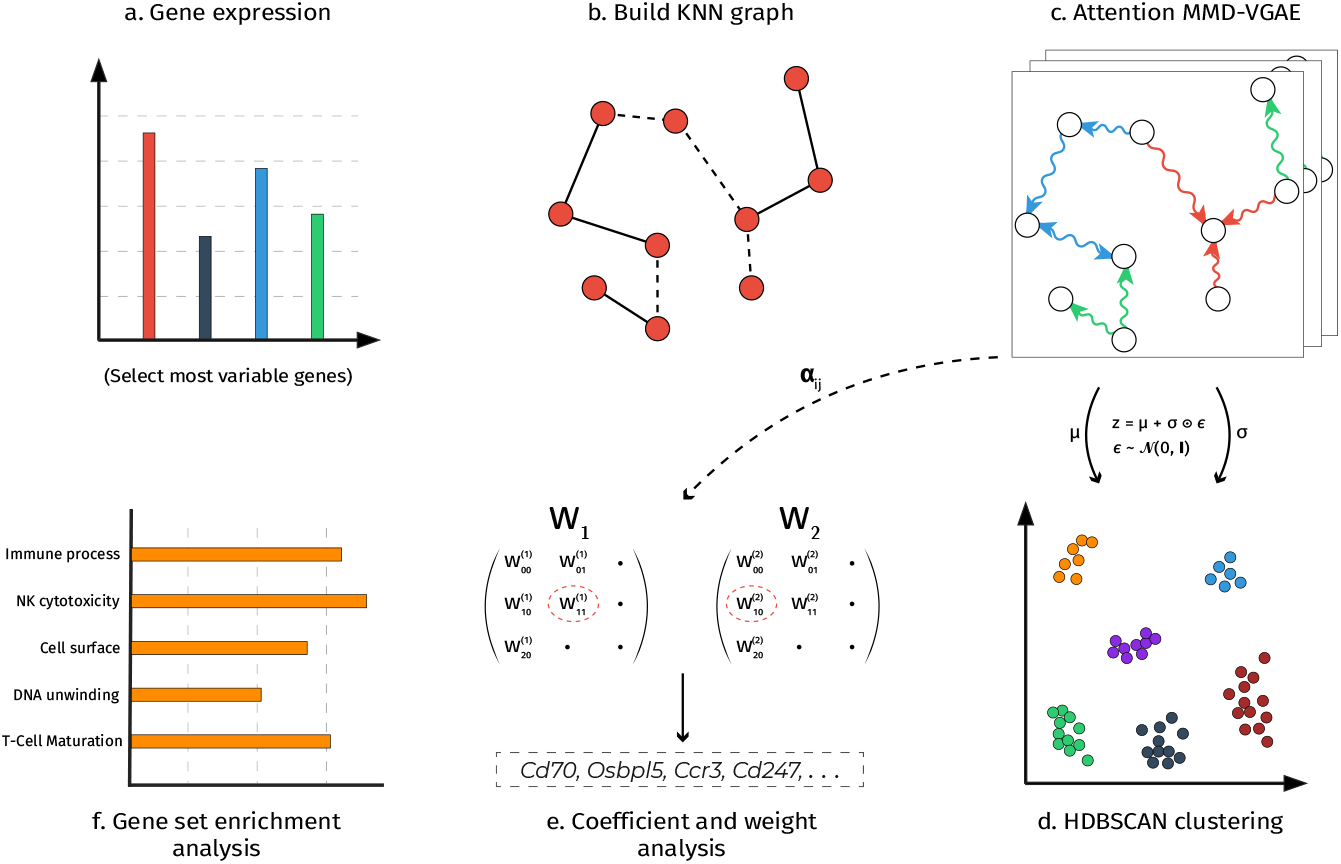
Simplified overview of the CellVGAE workflow. **(a)** After quality control and normalisation, the highly variable genes are selected. **(b)** A graph of cells is generated, in this case a *k*-nearest neighbour graph. **(c)** The CellVGAE model is trained using the graph structure and highly variable genes as node features. **(d)** The learnt cell representations are projected in 2D using UMAP and clustering can be performed, for example using the HDBSCAN algorithm. **(e)** Layer weight analysis reveals the most important genes in each cluster. **(f)** Gene set enrichment analysis is performed on the genes found in the previous step.

The main steps are:

1. **Feature selection** – We start with the usual quality control and normalisation steps that are present in all bioinformatics pipelines (we consider Seurat as a representative). CellVGAE requires log-normalisation to be applied to the count matrices (that have passed quality control). We successfully applied CellVGAE to different sequencing protocols and normalisation procedures that precede the log-transform (the 9 datasets in Section 5.3). For more details on these operations, we refer the readers to [LT19]. We use the standard in feature selection, namely selecting the most highly variable genes (HGVs). In our experiments we use either 250 or 500 HVGs, which become the node features.
2. **Graph generation** – This step involves the generation of connections between cells, i.e. a graph of cells. In this study, we mainly experiment with KNN graphs built using Euclidean distance in the original high dimensional space, alternatively using PCA dimensionality reduction before the KNN procedure for some datasets. The number of neighbours we have experimented with are *k* ∈ {5, 10, 20} as these include the default value of SAM (20), while the others are also recommended in the SAM documentation and we have observed better performance with smaller values of *k*. For two datasets (*Baron1* and *Segerstolpe*) we use the *k*^th^ closest cells given by the Pearson correlation on the selected features matrix. Although we are not aware of such existing techniques at the moment, in theory CellVGAE can also use a graph where the edges between the cells are experimentally (not computationally) derived. We have chosen KNN and Pearson graphs for simplicity, efficiency and their use in previous literature, but this does not restrict, in principle, the use of CellVGAE with graphs originating from other, perhaps more sophisticated algorithms.
3. **Training** – This step involves training CellVGAE using the data from the previous steps. The only additional preprocessing is in the form of min-max scaling applied to the HVG features. As a general recipe, we recommend using the same configuration of CellVGAE as we have used in Section 5. We recognise that an inherent difficulty in evaluating machine learning models on metrics such as the ARI or silhouette coefficient is that they are not used and monitored in the learning and validation steps. Concretely, only the loss from definition (3) can be monitored, which does not necessarily correlate with improved clustering performance. Our detailed strategy is given in Appendix C – Experimental Design.
4. **Projection and clustering** – The neural networks can now be used to find latent embeddings for the nodes (cells). Clustering directly on the learnt representations, even with advanced algorithms like HDBSCAN [MHA17], introduces noise, i.e. samples which are not able to be classified correctly. To circumvent this, we perform clustering using HDBSCAN on the UMAP 2D projection of the node embeddings. Due to the quality of the embeddings learnt by CellVGAE (Section 5), clusters are clearly delimited and separated, so for small datasets automatic clustering might not even be necessary.
5. **Interpreting the learnt representations** – Apart from the clustering itself, we can perform at least three different investigations, which will be fully detailed in the upcoming sections. These can be summarised as:

- Identifying the most relevant latent dimensions for each cluster and correlating them to the high-weight genes from the weight matrices. This is the same analysis from DiffVAE, but adapted to work with graph neural networks; effectively, we are not losing any features already introduced by DiffVAE.
- On a similar principle to the above, we can approximately determine which latent dimensions encode the most information for each top gene. This allows us to map expression vectors to the 2D cell projections, visualisable as a heatmap. Although approximate in nature, we are able to derive the expected results for more than 16 genes (8 examples given in Section 5.2.2). To the best of our knowledge, this is the first attempt using neural networks.
- Visualising the attention coefficients as a graph, indicating the contribution of each cell to its neighbours. This can also be extended to a mapping of the coefficients to the 2D projection of cells. We believe that analysing these learnt parameters can capture biological trends and interactions, which can be further used for generating hypotheses and follow-up studies. The possibility of using attention coefficients in this manner is exclusive to graph attention networks, and as such, in the context of scRNA-seq, to CellVGAE.
6. **Gene set analysis** – This is a possible step after identifying the most important genes for each cluster. For Figure 1, we used g:Profiler [RKK^+^19]. Another popular choice is Enrichr [CTK^+^13], [KJR^+^16].

## 5 Results

We begin our evaluation of CellVGAE by presenting three different exploratory study cases (one in Appendix A). The point of these experiments is not to evaluate the performance numerically (see Section 5.3), but to explore and demonstrate the capability of a graph neural network approach on real world datasets. All 2D visualisations in our study use UMAP.

### 5.1 The *Schistosoma mansoni* dataset

One of the motivating factors behind the SAM algorithm (Section 2.2) is the inability of the existing methods to analyse a novel scRNA-seq dataset of *Schistosoma mansoni* (including cell types such as *ϵ, δ*′ and *μ*). More specifically, Tarashansky et al. [TXL^+^19] show that commonly used methods like PCA, Seurat and SIMLR [WZP^+^17] fail to distinguish any cluster formations. In turn, this renders tasks like cell (sub)type identification and finding marker genes difficult using current tools. To formalise this notion of difficulty, the authors also introduce an unsupervised metric called *network sensitivity*, which measures the changes in cell-to-cell distances on randomly selected subsets of the gene expression matrix. An intuitive explanation of a high sensitivity score is that changes in the used features lead to a significantly different topological network. On the other hand, datasets robust to the said feature selection have many genes (signals) reinforcing the same structure, thus resulting in a low sensitivity score.

Out of the 56 datasets considered by the SAM authors, the *Schistosoma mansoni* one has the highest sensitivity score by a considerable margin (over 0.25 higher sensitivity score than the following dataset), i.e. it is the most difficult to analyse. Therefore, we consider the performance of CellVGAE on this dataset to be illustrative of the potential that graph-based neural methods have. The 9 datasets examined in Section 5.3.1 cover a lower range on the network sensitivity spectrum and are thus appropriate settings for studying the broader applicability of our method. The exact sensitivity ranking and numerical values are available in [TXL^+^19].

#### 5.1.1 Clustering

As a first step, we train CellVGAE on the preprocessed and log-normalised 250 top HVGs from the *Schistosoma mansoni* dataset and a KNN graph with *k* = 20, with 50 latent dimensions. We also train DiffVAE on the same HVGs dataset with default settings (roughly equivalent) and 50 latent dimensions, and Seurat by running the functions RunPCA and RunUMAP with 10 dimensions. We include the plot from SAM with default settings (as the documentation advises for this dataset) for comparison. We do not illustrate other methods as SAM already includes an exhaustive evaluation against the most widely used tools.

It is easily noticeable from Figure 2 that the DiffVAE and Seurat representations do not distinguish any clusters and make the application of density-based clustering impossible. On the other hand, CellVGAE finds three clearly delimited clusters, just as SAM, and moreover, we observe 95.30% overlap for the red cluster (142 common cells out of 149), 94.62% overlap for the blue cluster (88 common cells out of 93) and 76.04% overlap for the green cluster (73 common cells out of 96), where the overlap percent is calculated as the number of common cells over the number of cells in the original SAM cluster. To summarise, this leads to an average of 88.65% overlap between the clusters and an ARI of 0.7075, which is remarkable considering the small number of cells (total 338) and sensitivity of the ARI. CellVGAE is also only using the 250 most variable genes and a naive KNN graph, while SAM uses a bespoke algorithm that specifically addresses limitations of traditional feature selection and graph generation techniques. We note that for Seurat, it is possible to use Louvain clustering on an SNN graph built from the PCA representations (not the UMAP, as above), which can lead to a more meaningful clustering. For a detailed comparison of existing methods on this dataset, we refer the readers to the SAM study [TXL^+^19].

**Figure 2:**
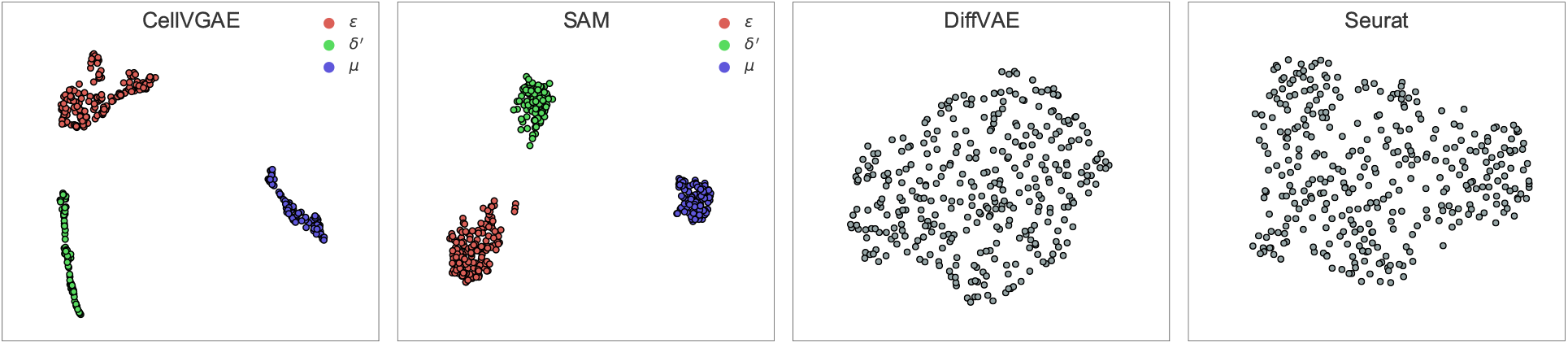
UMAP plot of the cell embeddings for four different methods. For CellVGAE and SAM, clusters with the highest overlap use the same colour in their respective plots.

#### 5.1.2 Finding marker genes

As mentioned in Section 4.1, by using graph attention layers we enable a very similar weights analysis procedure as in DiffVAE. A crucial difference is that DiffVAE uses a neural network decoder with multiple layers, while CellVGAE uses a simple inner product decoder. Still, weights analysis can be carried out on the encoder, thus linking the latent dimensions to the genes. Although the mechanism of graph attention layers is different from the simpler dense layers, each GAT layer is parameterised by a learnable weight matrix **W** (Section 3). Assuming a CellVGAE architecture with the same number of attention heads, two hidden layers: 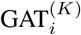 with *i* ∈ {1, 2}, and two layers for the parameters: 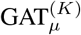 and 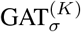, where *d_i_* is the output dimension of each layer, we can extract the weight matrices 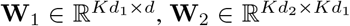 and 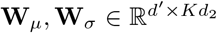, where additionally *d* is the input number of dimensions, *d*′ is the number of latent dimensions and the matrix dimensions follow PyTorch conventions. These allow us to derive the following products for the ***μ*** and ***σ*** layers:

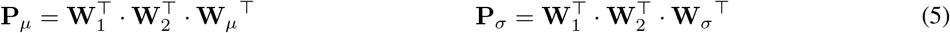

with dimensions *d* × *d*′. These matrices effectively link the genes to the latent dimensions. We can now apply the DiffVAE procedure described in Section 2.1. For each cluster, and then for each latent dimension, the 15 top genes are selected. Overall, the same gene might be selected multiple times, so we count the number of appearances of genes for all clusters. We report some of the found genes in descending number of appearances in Table 1 and additionally, if one gene appears only one time, but is within the top 100 weights in the product matrices, we also include it in the bottom row.

Out of the genes mentioned in the SAM manuscript, only 5 are in the top 250 variable genes, and CellVGAE correctly identifies them. More specifically, the RNA binding protein *nanos-2* (identifier 051920), which is characteristically expressed in *δ*′ cells, *eled* (identifier 041540) which marks *E* cells, an actin protein (identifier 161920) and *dhand*, a helix-loop-helix transcription factor (identifier 062490) that mark *μ* cells and finally an aschaete-scute transcription factor, *astf* (identifier 142120) which is highly enriched in *ϵ_α_* cells, a subpopulation of *ϵ* cells. We notice that CellVGAE cannot distinguish between the two subpopulations of *ϵ_α_* and *ϵ_β_*, but the gene that is abundantly expressed in *ϵ_β_* cells, (identifier 087310, codes for *bhlh*) is not even present in the 250 HVGs, among others in the same situation.

**Table 1:** (Top row): Some high-weight genes found by CellVGAE in descending order. These genes appear multiple times across latent dimensions (more than 2). (Bottom row): Some genes that appear once but have high values in **P**_*μ*_. Bold genes are also identified as marker genes by SAM.

| Matched cluster | $\epsilon$ (red) | $\delta'$ (green) | $\mu$ (blue) |
| --- | --- | --- | --- |
| Latent analysis | 129510, 191690, **070380**<br>**161920, 021340** | **161920, 070380**, 179790,<br>**041550, 068280** | **068280**, 169030, **105220**,<br>**161920, 070380, 041550** |
| Top 100 | **142120, 041540** | **051920, 158740** | **062490, 158740, 044680** |

### 5.2 The PBMC3k dataset

We now consider a well-known dataset, PBMC3k^1^, of 2638 cells (about 8 times larger than the *Schistosoma mansoni* dataset). We use a very similar setup as in Section 5.1, including a dataset of the top 250 HVGs, with the same CellVGAE architecture, only with a reduction in attention heads from 20 to 16 for memory saving purposes and an increase in dropout.

#### 5.2.1 Clustering performance

Although PBMC3k is routinely analysed with established methods like Seurat^2^ or ScanPy^3^ [WAT18], we noticed that both SAM and DiffVAE struggle with this dataset. We benchmarked SAM extensively by performing separate runs with parameters `k` ∈ {5, 10, 20, 40, 80, 120, 160, 200} (number of nearest neighbours) and `num_norm_avg` ∈ {1, 5, 10, 25, 50, 100, 150, 200, 800} (number of largest spatial dispersions to average), all values being provided in the official documentation, for a total of 72 configurations. None was able to find more than 5 clusters. DiffVAE was run 10 times with settings recommended by the authors in [BATCL20], no version succeeding in finding more than 3 clusters. The performance of SAM and respectively DiffVAE in terms of cluster overlap and ARI was extremely close across all trials. An illustration of the clustering performance (on the 2D UMAP representation) is given in Figure 3.

**Figure 3:**
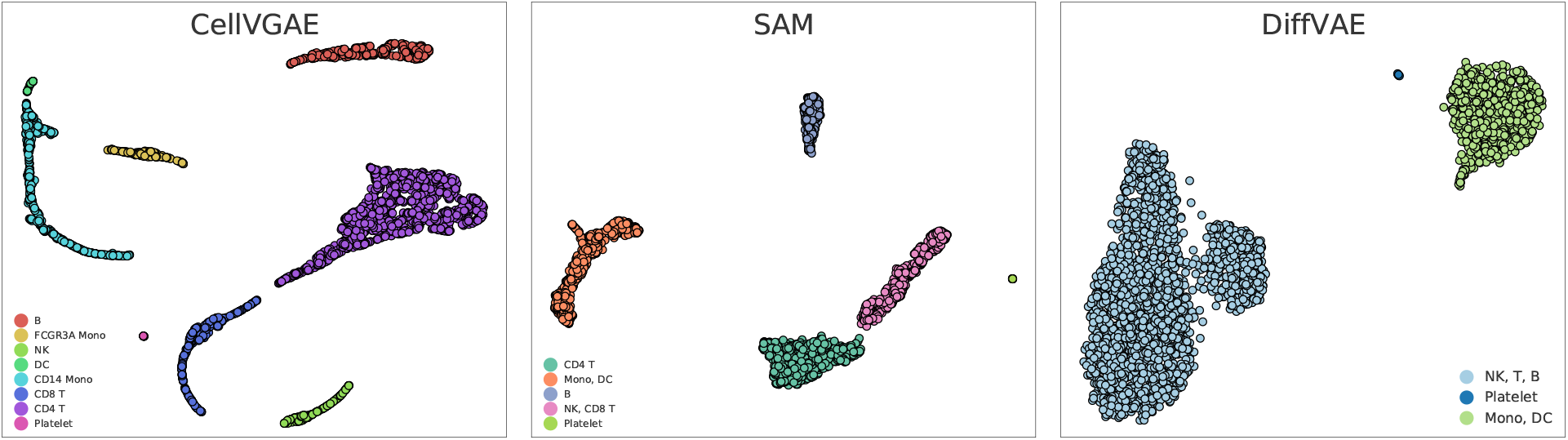
Clustering performance of CellVGAE, SAM and DiffVAE on the PBMC3k dataset.

For Figure 3, labels are determined by the highest overlap with Seurat clusters. CellVGAE shares 99.42% of B cells, 98.75% of CD14 Mono, and 96.27% of CD4 T cells with Seurat, with the others having an overlap of slightly over 87% each. The only exception is for Platelets at 78.57%, however this is due to the extremely small cluster size (CellVGAE correctly finds 11 out of 14 Platelet cells). Overall, CellVGAE achieves an ARI of 89.74% computed using Seurat as the “ground truth”, while SAM stands at 75.40% and DiffVAE at 37.60%. We recognise that Seurat is a computational method as well and does not represent the ground truth; however, CellVGAE is the only method we tested that can find the correct number of clusters, and with a high ARI, giving us confidence that it performs well in real-life situations. We also believe that an architecture tuned specifically for this dataset could achieve even better performance. These results suggest that while SAM uses the most sophisticated graph algorithm, the approach is not enough for correct classification on certain datasets. In this case, SAM cannot distinguish the NK population from the CD8 T population, both being relatively large, and neither the more finely-grained Monocytes and DC cell populations. On the other hand, when trained on the same dataset as CellVGAE, DiffVAE groups NK, CD4 T, CD8 T and B cells together in a single cluster. Similarly, Monocytes and DC are recognised as a single cluster like in SAM. These observations suggest that supporting neural network with graphs (like in CellVGAE) can outperform the individual methods taken in isolation.

#### 5.2.2 Visualising gene expression

Inspired by the analysis in Section 5.1.2, we make a first step towards visualising what the model has learnt in terms of gene expression. This type of analysis can suggest the contribution of a gene at the level of each cell, providing a granular view of expression within a cluster. For this purpose, we make use of the learnt node embeddings, a matrix **E** of dimension *N* × *d*′, where *N* is the number of cells and *d*′ is the latent (output) size, as well as of **P**_*μ*_ with dimension *d* × *d*′. We then simply select the vector **g** corresponding to a target gene of choice out of **P**_*μ*_ and perform the multiplication **E** · **g**, which gives the CellVGAE expression value, used as the hue parameter when plotting. This approximate procedure provides an intuitive visualisation of expression from the neural network’s perspective, and produces results in agreement with existing knowledge. In Figure 4 we present plots for 8 different genes from the 250 HVGs we used. We chose to interpret positive values as a form of high expression, and negative values as the opposite.

**Figure 4:**
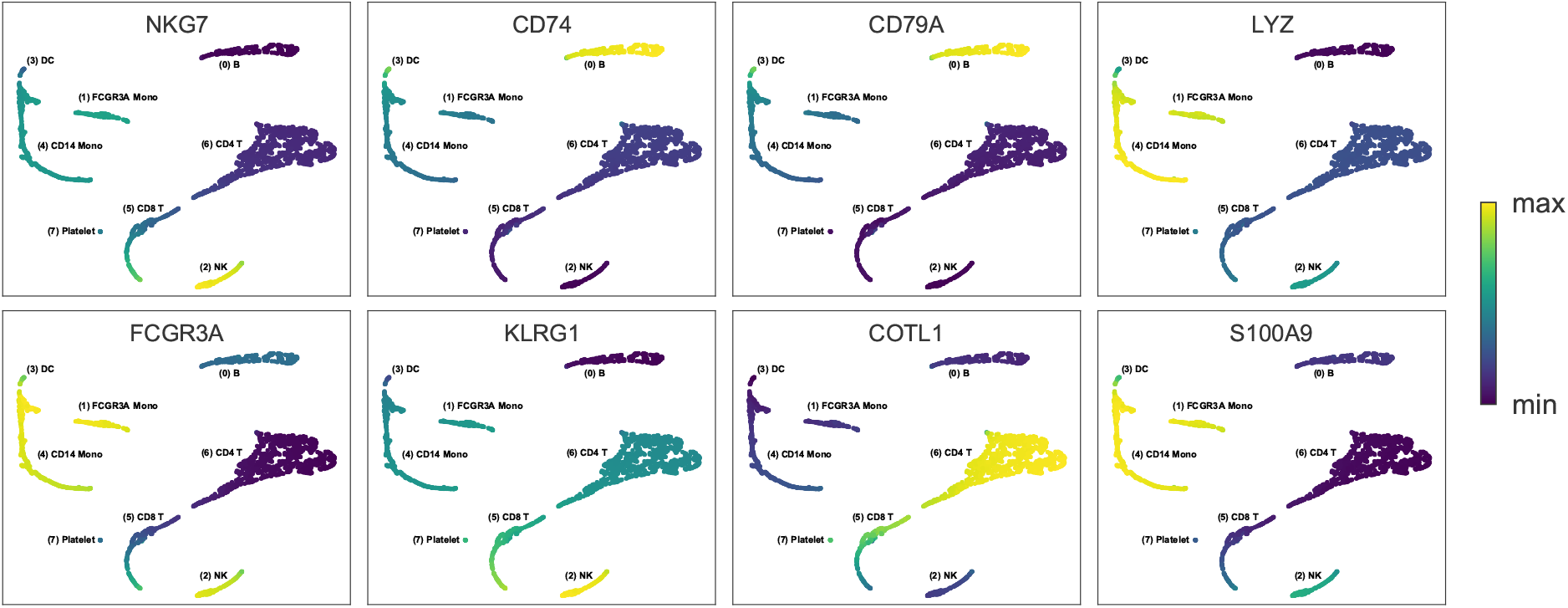
Visualisation of CellVGAE expression for 8 different genes from the HVG PBMC3k dataset. Each plot uses a different scale.

### 5.3 Evaluating performance on cell clustering

Having discussed the exploratory potential of our model, in this section we provide an extensive evaluation of CellVGAE, SAM and DiffVAE on 9 well-annotated scRNA-seq benchmarks where we show improved clustering performance. Information about the datasets is summarised in Appendix B – Table 3.

#### 5.3.1 Results

Our decision to use these 9 datasets is not by chance; with only one exception which will be clarified shortly, these are all datasets that were used to benchmark SAM in [TXL^+^19]. We run SAM ourselves for all experiments, as we report additional metrics, but the utilisation of these datasets as benchmarks in previous work increases our confidence in choosing appropriate evaluation strategies and in the results that we obtain. The one exception is the dataset *Loh* – where SAM results are reported on a version of the dataset with 651 cells; however, while we were able to find this version on the conquer database, it has over 65,000 rows (genes) and we have discovered that the standard approach of selecting the top HVGs does not work on this massive dataset, i.e. it leads to untrainable neural models with very poor performance (both CellVGAE and DiffVAE). For this reason, we have replaced the dataset with a filtered version of 498 cells and 20,142 rows, provided by the original publication authors, amenable to HVG selection, but we acknowledge that SAM works successfully even on the 651 cells version. We also note that the DiffVAE authors have already benchmarked the method on a different version of the *Muraro* dataset, with 2285 cells, hence the results are not comparable. We were unable to find this version online, and since SAM uses the version with 2126 cells, we use it as well, for both CellVGAE and DiffVAE.

By examining Table 2, we observe that CellVGAE obtains the highest ARI score on every dataset, some being very close (e.g. on *Muraro*, *Baron1*), while others, such as *Darmanis*, *Wang*, *Baron4* are at a large distance (at least 3 ARI points). We notice that DiffVAE performs better on the larger rather than smaller datasets, which is expected for neural networks. We attribute the success of CellVGAE even on relatively small datasets to the underlying graph representation, an additional structure compared to DiffVAE that is exploited successfully during the learning process.

**Table 2:** Clustering results on the 9 datasets from Table 3. The metrics used are the adjusted rand score (**ARI**), silhouette coefficient (**SC**), number of clusters found (**#**) and noise (**N**). Noise represents the number of cells which cannot be assigned to a cluster according to the clustering algorithm. For CellVGAE, in certain datasets with many small clusters, i.e. *Baron1*, *Baron2*, it is possible to separate additional small clusters manually (missed by HDBSCAN) for additional performance.

|  | Truth | SAM |  |  | CellVGAE |  |  |  | DiffVAE |  |  |  |
| --- | --- | --- | --- | --- | --- | --- | --- | --- | --- | --- | --- | --- |
|  | # | ARI | SC | # | ARI | SC | # | N | ARI | SC | # | N |
| Darmanis | 8 | 0.9112 | 0.6966 | 8 | **0.9472** | 0.7724 | 8 | 0 | 0.8621 | 0.7104 | 7 | 4 |
| Wang | 7 | 0.8986 | 0.2794 | 7 | **0.9418** | 0.4772 | 5 | 1 | 0.9213 | 0.5939 | 5 | 5 |
| Baron1 | 14 | 0.9644 | 0.7314 | 11 | **0.9670** | 0.7476 | 11 | 0 | 0.9659 | 0.6953 | 10 | 27 |
| Baron2 | 14 | 0.9680 | 0.6726 | 10 | **0.9817** | 0.7139 | 9 | 0 | 0.9428 | 0.6926 | 8 | 6 |
| Baron3 | 14 | 0.9460 | 0.6839 | 8 | **0.9723** | 0.7376 | 10 | 0 | 0.9459 | 0.6626 | 8 | 56 |
| Baron4 | 14 | 0.9263 | 0.7823 | 8 | **0.9764** | 0.7474 | 7 | 0 | 0.9287 | 0.7849 | 7 | 0 |
| Loh | 9 | 0.9637 | 0.7419 | 9 | **0.9684** | 0.8187 | 9 | 0 | 0.9560 | 0.7503 | 9 | 8 |
| Segerstolpe | 14 | 0.9356 | 0.3866 | 15 | **0.9734** | 0.6834 | 10 | 0 | 0.9667 | 0.7019 | 9 | 0 |
| Muraro | 10 | 0.9254 | 0.7303 | 9 | **0.9503** | 0.7272 | 8 | 0 | 0.9496 | 0.8556 | 8 | 0 |

We also include the silhouette coefficient (SC) in our reported results. Although we prioritise the ARI as a metric of clustering performance, the SC is a popular metric that, briefly speaking, can give a good indication of how well an object was assigned to its cluster. We note that generally all methods achieve silhouette scores that can be considered good (≥ 0.65), a few exceptions being some very low scores for SAM. The silhouette coefficient penalises clusters that are more spread out or not clearly delimited. To better understand CellVGAE’s silhouette scores, we provide an illustration of the clustering on the *Muraro* dataset in Figure 5, as it has one of the lowest SC scores relative to the other methods (also see Figure 9 in Appendix D). Notice that CellVGAE tends to find more complicated shapes (with holes, arcs, etc.), for example the Beta cells resemble a “Y” shape, while DiffVAE favours circular shapes and SAM generally contiguous shapes. These observations also apply to Figure 2 and Figure 3. Additionally, some CellVGAE clusters exhibit a linear shape. While we are currently investigating this, we suspect that the linearity and non-circularity of some clusters might be indicative of differentiation trajectories or other changes in cell properties.

**Figure 5:**
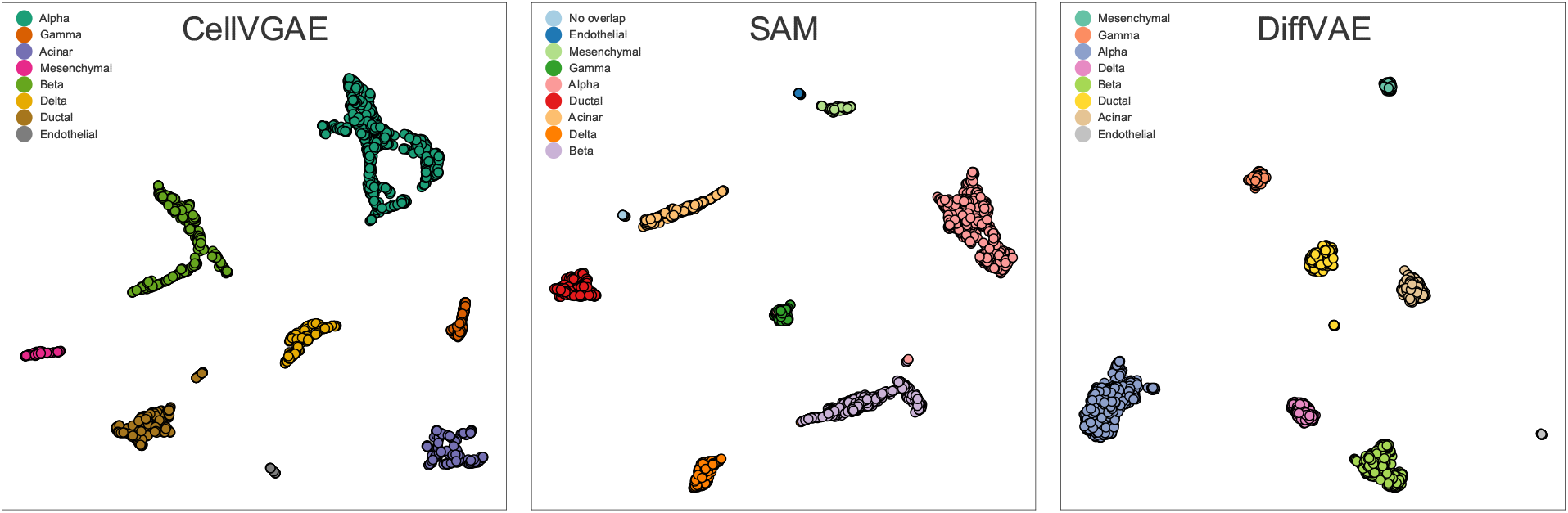
Different shape tendencies on the *Muraro* dataset.

In Table 2 we also keep track of the number of clusters found by each method (**#**). We observe that none of the methods is consistent in finding the true number of clusters, but arguably SAM is the closest. However, since this isn’t reflected in an increase in SAM’s ARI, we cannot say that it is meaningful. On the other hand, many of the clusters are extremely small, for example *Baron4* has 1 epsilon cell, 1 mast, 1 Schwann, 1 T, 2 acinar, 5 quiescent stellate and 7 endothelial cells. Realistically, we would not expect most methods, especially deep learning, to detect these clusters, although CellVGAE manages to find and separate the 10 macrophage cells (extreme left in the leftmost plot in Figure 9).

Finally, for both neural models we report the number of cells that were not clustered by HDBSCAN, which we call noise (N). This does not apply to SAM as it internally applies a further clustering step to HDBSCAN outliers using kNN classification. Complete details of their implementation are given in the manuscript [TXL^+^19] and source code. We do not apply further clustering, but we employ a postprocessing step to ensure that very small clusters are not missed by HDBSCAN. Our own procedure is detailed in Experimental Design – Appendix C. We notice that DiffVAE is more susceptible to noise, as only 3 out of 9 benchmarks produce noise-free representations. However, we do not expect the lower noise values (< 5) to be problematic. CellVGAE produces higher-quality embeddings in this regard, with a maximum noise value of 1 occurring only once.

### 5.4 Interpretability

In this section, we propose two alternative ways of visualising the learnt attention coefficients (denoted by *α*). To quickly recapitulate, the attention coefficients measure the contribution of each cell to its neighbours in a pairwise manner. Thus, the numerical value associated with two nodes can be interpreted as an edge. Note that for two nodes *i*, *j* the coefficients *α_ij_*, *α_ji_* are different, resulting in a graph which is weighted *and* directed. Each graph attention layer has its own independent set of coefficients. We emphasise that this type of analysis is unique to graph attention networks and, in the context of scRNA-seq, to CellVGAE.

Firstly, to provide a broad picture of the attention activations, we propose a visualisation where they are mapped to the 2D UMAP projection. Since the coefficients represent edges between vertices, to get a node-level representation we simply average over each node’s neighbourhood. The highest resulting values will give the nodes with the largest contribution in the neural model. Secondly, a finer-grained visualisation is to simply plot a graph of the nodes, where the *α* coefficients give the edge weights. A simple strategy is to select the top *k* largest coefficients; alternatively and depending on the specific use case, the analysis can exclude certain cell types or focus on a different range of interactions (e.g. the smaller edge weights).

We put the above methods in practice and give illustrations of the results in Figure 6 and Figure 7 for two datasets. We have chosen the *Darmanis* dataset since there is a relatively large difference in ARI compared to the other two methods of Table 2, so insights into the model are valuable. We also include the PBMC3k dataset since we have discussed it at length in the previous sections. For both figures and datasets, we chose the second layer of the CellVGAE model and for simplicity have taken the mean across all the attention heads, but the same analysis applies per-head and can, in fact, reveal the differences in what each one learns. The values are normalised with `matplotlib.colors.Normalize` to the shown range.

**Figure 6:**
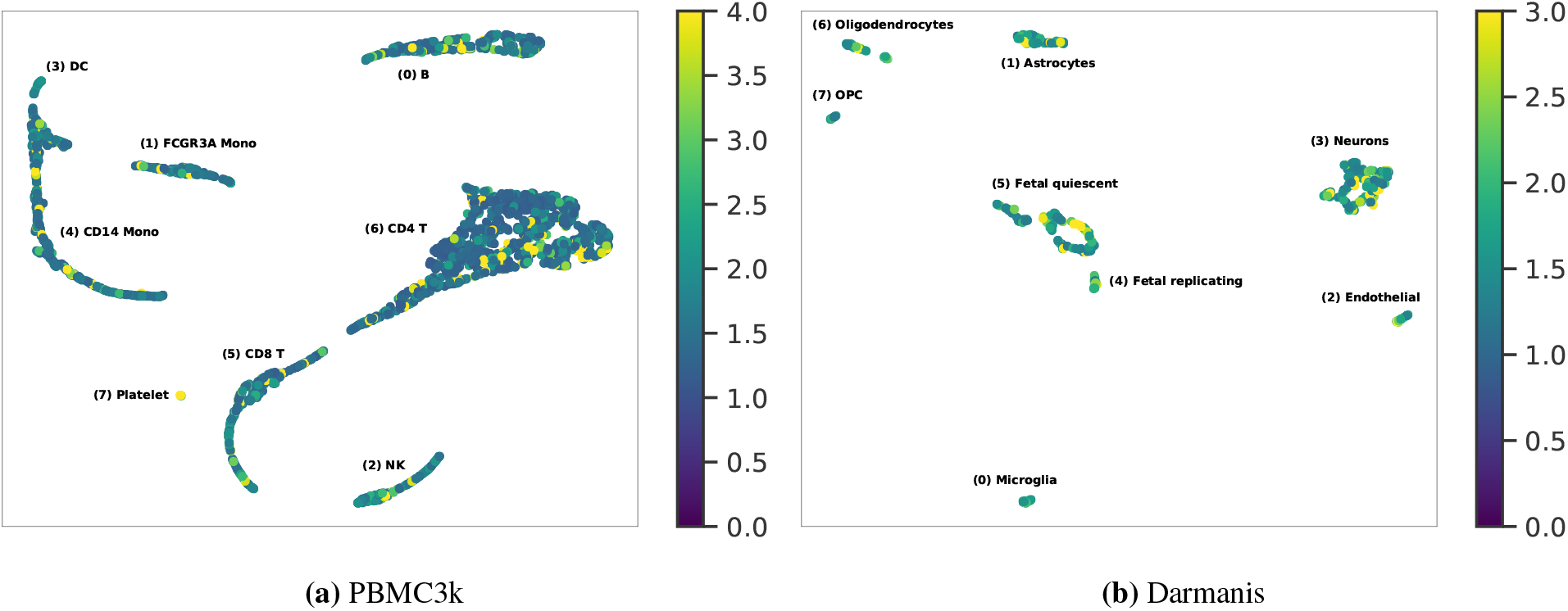
Attention coefficients, averaged across all heads and normalised, mapped to the 2D UMAP projection.

**Figure 7:**
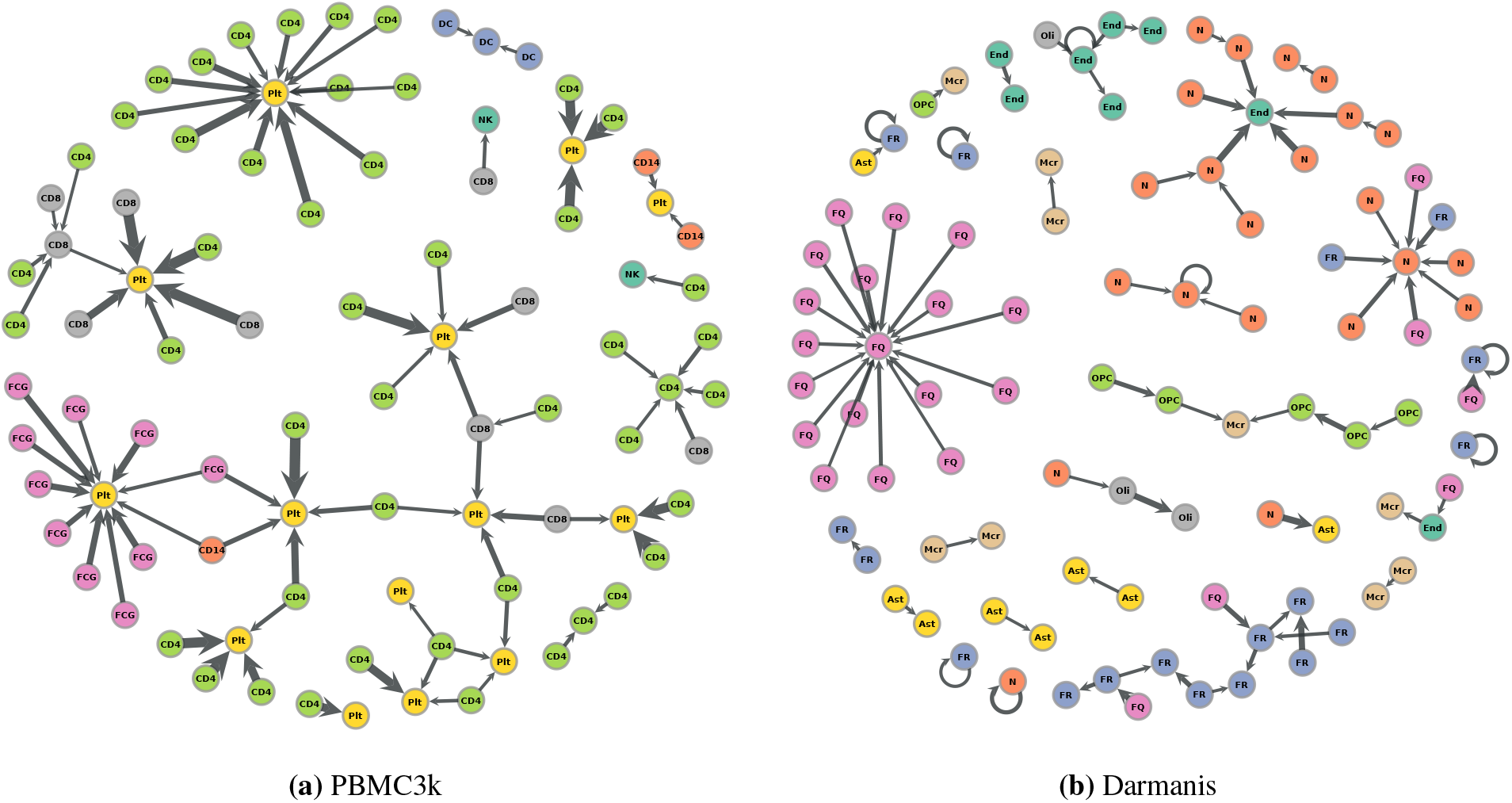
Weighted graph given by attention coefficients (top 80). Cell names are abbreviated and the list of full names is provided at the end of the section. Edge widths use different scales. Plots generated using graph-tool. [Pei14].

For PBMC3k, a surprising finding from the UMAP plot in Figure 6 is that the Platelet cells are highly involved at the level of the second layer. In fact, by inspecting the graph representation, we discover that all 14 Platelet cells are among the top contributors, with some of the highest-weight edges (attention coefficients). The surprise lies in the fact that the Platelet cluster is by far the smallest, at just over 0.005% of all cells. This interesting dynamic could be a point of further biological study. It is known in the literature that platelets can regulate immunitary functions and interact with other immune cells ([LVR15], [GSH^+^15], [SCRE09], [PWPW20]). These interactions might be indicative of an underlying condition or capture a particular phenomenon that can support a biological hypothesis. Otherwise, we notice that the CD4 T and CD8 T cells dominate the top 80 plot, and this can be probably attributed to their high number. Another prominent group is given by the FCGR3A cells. For this particular graph, we notice the existence of long paths and a relatively low number of connected components.

For the *Darmanis* dataset, all 8 cell types (see below) are encountered, suggesting that multiple different cell types can inform the classification of a single cell, as can be seen in the relatively heterogeneous graph in Figure 7 (b). Neurons, which are among the most activated, can be seen interacting with an Endothelial cell. Another large group, of Fetal quiescent cells, is seen largely interacting with its own kind, although it participates in other relationships, mostly with Fetal replicating cells. On this dataset, we notice the presence of self-loops. These might indicate that the cell’s own features are enough to distinguish itself. Overall, the graph is more fragmented (more connected components) indicating less cross-talk.

The cell types illustrated in Figure 7 (a) are: Platelet (Plt), CD4+ T (CD4), CD8+ T (CD8), FCGR3A+ Monocyte (FCG), CD14+ Monocyte (CD14), Natural Killer (NK) and Dendritic Cell (DC). For Figure 7 (b): Fetal quiescent (FQ), Fetal replicating (FR), Astrocyte (Ast), Neuron (N), Endothelial (End), Oligodendrocytes (Oli), Microglia (Mcr), OPC (OPC).

## 6 Conclusion

We have introduced CellVGAE, a machine learning architecture that integrates the capability of graph techniques with recent advancements in neural networks. Starting with a general workflow, we have explored the potential and applicability of CellVGAE in a number of different, carefully chosen scenarios. We have shown that our method performs consistently well in finding accurate, informative clusters, even when applied to complex datasets with subtle signals or when existing state-of-the-art methods perform poorly. Furthermore, we extensively evaluated CellVGAE on 9 well-annotated datasets of different sizes and complexities, using the most popular and relevant clustering metrics. The high scores achieved throughout all benchmarks provide evidence of the architecture’s power and wide applicability. Interpretability is a first-class consideration when designing CellVGAE, and we have delivered three strategies: (1) high-weight gene identification, useful for gene markers and gene set enrichment analysis; (2) visualisation of learnt expression, per-gene and (3) a practical interpretation of the attention coefficients.

### 6.1 Limitations and future directions

Having presented and described our model in detail, we now turn our attention to possible limitations. We identify two main ways in which the architecture could be expanded upon. Firstly, all the graphs we use are based on simple measures, for example on the Euclidean distance in the high-dimensional space. As we have shown, this approach clearly has value, as it performs consistently well in all of our experiments. It is conceivable that different distance metrics could be used, perhaps depending on the dataset, or when similarity is not the main concern. After we have introduced this scRNA-seq analysis workflow, bespoke graph creation algorithms might be designed in the future, specifically for deep learning. We believe that we have made the first steps in this direction and leave these suggestions for future study.

Secondly, we subscribe to the idea of using a relatively small number of gene features per node, in order to keep the node dimensionality smaller than the number of nodes in the graph. It is not generally expected that message passing will perform as well with very large node features, and this could be desirable for some particular datasets. As graph convolution operators are an active research area now, it is possible that in the future this drawback will no longer exist.

Furthermore, although we have focused on scRNA-seq data, it should be possible to adapt the architecture to other biological data types, such as ATAC-seq, where the input would be the peak count matrices. On this note, integration of multiple modalities seems within reach, as autoencoders are especially suited for the task. Finally, as machine learning and graph neural networks in particular are rapidly advancing research topics, future innovations can be integrated into CellVGAE. These include but are not limited to different graph convolutions and different regularisation losses (for example, the recent Geometric Jensen-Shannon Divergence [DSL20] for variational autoencoders). In principle, all methods provided by the PyTorch Geometric library are accessible. A potentially interesting approach is graph pooling, where the graph topology can be downsampled. The obvious application is to improve performance, but the pooling could also be applied to compress the graph, with possible applications in clustering and trajectory inference. On the same note, a further interesting study might be in relating the CellVGAE cluster shapes to potential cell trajectories or other such properties.

## Appendices

## Appendix A The macrophages dataset

Lane et al. [LVVD^+^17] introduced a scRNA-seq dataset to study dynamic patterns of NF-kB activation. According to the network sensitivity introduced by SAM, the macrophages dataset is the second most challenging dataset, after the *Schistosoma mansoni* dataset, with a numerical value of just under 0.4 sensitivity. We follow the same strategy of comparing the clustering results as in Section 5.1 and the same parameters. We do not include Seurat or SIMLR [WZP^+^17], as these are discussed in the SAM manuscript. An illustration is provided in Figure 8.

**Figure 8:**
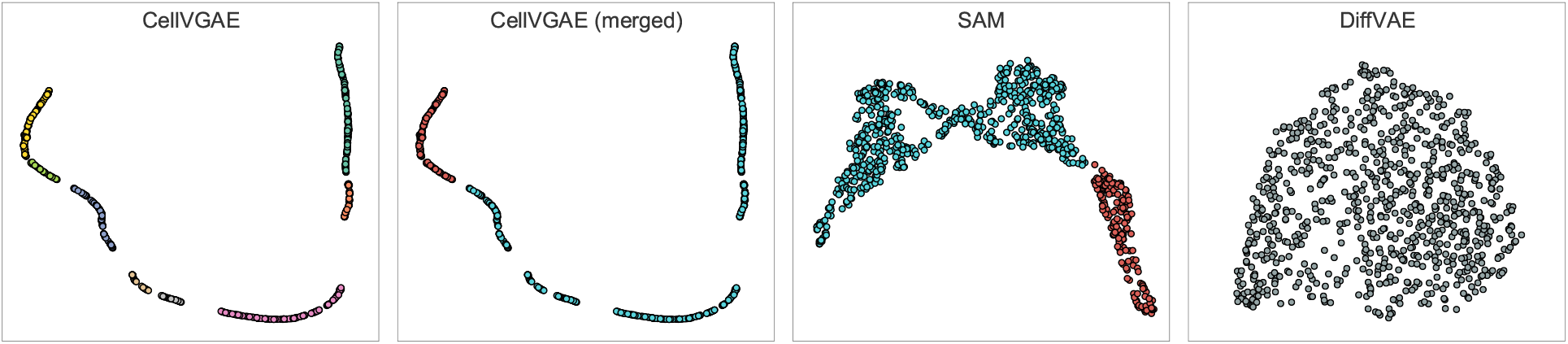
Clustering performance of CellVGAE, SAM and DiffVAE on the macrophages dataset.

CellVGAE finds a larger number of clusters, depending on the granularity of HDBSCAN (pictured are 8 cell formations). SAM finds two clusters, as reported in the study, without providing much information about their identity. We notice the lack of a clear boundary between their clusters. The SAM authors then proceed to study the dataset from an activation dynamics perspective, by removing the cell cycle effects. Here, we focus on the transcriptomics. We find our results interesting for two reasons: (1) the clusters can be merged to recover the SAM clusters, with 95.00% overlap with the large (blue) cluster (608 out of 640) and 91.26% overlap for the smaller (red) cluster (167 out of 183), for an average overlap of 93.13% and an ARI of 0.7604; (2) Lane et al. report multiple clusters (10) based on the transcriptomics data in Table S3. Unfortunately, their dataset clusters are not available. We consider the existence of multiple clusters and the ability to largely recover the SAM clusters encouraging. DiffVAE is unable to distinguish any clusters.

## Appendix B Datasets

We collect and label the datasets used in Section 5.3.1, along with the accompanying publication, size and source of download in Table 3.

**Table 3:** Summary of the benchmark datasets.

| Name | Publication | # of cells | Source |
| --- | --- | --- | --- |
| Darmanis | [DSZ<sup>+</sup>15] | 420 | single-cell-sota |
| Wang | [WSW<sup>+</sup>16] | 457 | hemberg-lab |
| Baron1 | [BVW<sup>+</sup>16] | 1937 | Gene Expression Omnibus, supplementary file |
| Baron2 | [BVW<sup>+</sup>16] | 1724 | Gene Expression Omnibus, supplementary file |
| Baron3 | [BVW<sup>+</sup>16] | 3605 | Gene Expression Omnibus, supplementary file |
| Baron4 | [BVW<sup>+</sup>16] | 1303 | Gene Expression Omnibus, supplementary file |
| Loh | [LCK<sup>+</sup>16] | 498 | Cell, supplementary file |
| Segerstolpe | [SPE<sup>+</sup>16] | 2209 | hemberg-lab |
| Muraro | [MDG<sup>+</sup>16] | 2126 | hemberg-lab |

## Appendix C Experimental design

As we aim to fairly and comprehensively evaluate the three methods, we start by describing our experimental design.

## C.1 Dataset preprocessing

We assume the data has already passed quality control (except for the *Wang* dataset, which comes with 178 low-quality samples that are discarded by us and by SAM). For SAM, we input the count matrix as-is, since the algorithm has its own preprocessing method and works on the entire gene expression matrix. For CellVGAE and DiffVAE, we log-normalise the data and select the most variable genes using Seurat (`NormalizeData` and `FindVariableFeatures` functions). Our philosophy is to select the minimum number of variable genes that still allow for state-of-the-art performance. There are two immediate benefits for this approach. Firstly, it is computationally more efficient and more appropriate for graph machine learning, since it is regarded as beneficial to keep the number of features per node smaller than the number of nodes. Secondly, by using only a restricted subset of genes and achieving leading performance, we are assured that the genes in the subset are sufficient to learn quality representations, the other genes being redundant. Since a few of the smaller datasets revolve around 500 cells in size, we chose to experiment with values of 250 HVGs (half the dataset size) or 500 HVGs (for larger datasets). DiffVAE uses the same datasets as CellVGAE, only without the graphs. Since DiffVAE uses dense layers, we acknowledge that it might prefer datasets with a large number of genes (e.g. 1500 or more). However, the DiffVAE authors successfully train the model with datasets of 700 genes and in our own evaluation it is often able to outperform SAM even on the smaller datasets.

## C.2 Graph generation

Our strategy for graph generation in the 9 experiments is to start with the *k* = 5 KNN graph built from the selected HVGs with the scran [LMM16] function buildKNNGraph. Slight variations are possible and were found to increase performance, for example in the form of applying *d*-dimensional PCA before the nearest neighbour search. This is applied by specifying a d parameter to the buildKNNGraph function. For the *Darmanis* dataset, building the KNN graph from the entire log-normalised gene expression matrix, while also applying PCA helped (the node features are still the 250 HVGs). Alternatively, for two datasets we use the Pearson correlation graph (each cell is connected to *k* other cells with the highest correlation values). Detailed information about the setup used for each dataset is provided in the repository.

## C.3 Architecture

Architecturally, we use a two-layer encoder with 128 hidden dimensions, 50 latent dimensions and the same settings of learning rate and batch size for all datasets, only lowering the number of attention heads for the larger datasets. We note that all experiments are performed on a consumer-grade mid-range GPU equipped with 8GB of VRAM and we did not hit any memory limitations when using the KL loss. A large number of attention heads is not required for top performance, but we have followed the indication of the GAT authors that increasing the number of attention heads can stabilise the learning. The default PyTorch implementation of the KL loss is considerably more memory efficient than the default MMD loss, and we recommend it for large datasets. DiffVAE parameters are based on the default, recommended settings and generally mirror CellVGAE’s, with two layers of sizes {256, 128} in the encoder and decoder, and 50 latent dimensions.

## C.4 Experimental pipeline and training

We have observed that running SAM with default parameters can lead to poor performance on a number of datasets. This is not unexpected as the datasets vary greatly in size, composition and scRNA-seq protocol. To ensure we show the best that the method is capable of, we have run SAM with the same selection of `k` ∈ {5, 10, 20, 40, 80, 120, 160, 200} and `num_norm_avg` ∈ {1, 5, 10, 25, 50, 100, 150, 200, 800}, as used in Section 5.2.1, and report only the best value.

For the neural approaches, a complication that arises is that the ARI cannot be optimised directly. To combat this, we first use a 90/10 split for each dataset and monitor the validation loss, to detect the point of convergence and then (eventually) overfitting. Once this is established, we propose a universal strategy where we train different models for 100, 120, 140, 160, 180, 200 and 250 epochs for DiffVAE, and 200, 220, 240, 260, 280, 300 and 400 epochs for CellVGAE, on the whole dataset. We train CellVGAE for more epochs as the convergence occurs later. We run each configuration 20 times, for a total of 140 trials for each dataset, or 1260 trials in total, for each of CellVGAE and DiffVAE. We make the observation that since both methods rely on clustering applied to the 2D UMAP projection of the latent embeddings, the UMAP representation itself is important. UMAP is not deterministic unless using the same seed and we have observed a broad range of resulting ARI scores depending on the parameters of UMAP. To filter out irrelevant representations, we run UMAP 30 times with random initialisation, for each trial. Finally, for each resulting projection, we run HDBSCAN 21 times with the min_cluster_size and min_samples parameters locked to the same value in {10, 11,…, 30}. Due to the large variability in data quantity and structure, a single clustering setting would be unable to perform well across all datasets and projections. As such, we remove any bias by using a large search space. After applying all the steps above, we select the best model in terms of ARI and report it, along with the silhouette coefficient, number of clusters found and noise, i.e. cells classified as “-1” by HDBSCAN (Table 2). While this evaluation procedure is involved, we have found that for purely unsupervised studies using CellVGAE, a high silhouette score (relative to the dataset) can be a good predictor of high ARI.

If cells are excluded due to HDBSCAN parameters (e.g. minimum cluster size too high for the very small clusters) but would otherwise form a well-defined cluster, they are not counted towards noise and contribute towards the # metric in Table 2. Thus, the cells included as noise are only those who are too close to existing clusters to be properly classified.

## C.5 Experimental platform

CellVGAE was developed using PyTorch [PGM^+^19] and PyTorch Geometric [FL19], at version 1.6.0 and 1.6.1 respectively (also tested successfully with version 1.7.0 of both). The other tools and libraries used the most recent version available for Python 3.8 at the time of writing. Most plots use matplotlib [Hun07] and seaborn [Wtsdt20]. Seurat 3 was installed on R version 4.0.2 [R C13]. The main development operating system was Windows 10, running the latest insider developer build available at the time of writing. Tools available exclusively on Linux were installed and used on Ubuntu 20.10 running on the same hardware. All CellVGAE models were trained on a NVIDIA GeForce RTX 2070 GPU with 8GB VRAM connected through Thunderbolt 3 to a machine equipped with an Intel Core i9-8950HK and 32GB of DDR4 memory. The used CUDA toolkit version is 10.2 for PyTorch 1.6.0 and 11.0 for PyTorch 1.7.0. DiffVAE was run exclusively on the CPU, using a compatible version of TensorFlow (1.15).

## Appendix D Clustering on the *Baron4* dataset

In addition to Figure 5, we provide an illustration of the clusters and their shape on the *Baron4* dataset in Figure 9.

**Figure 9:**
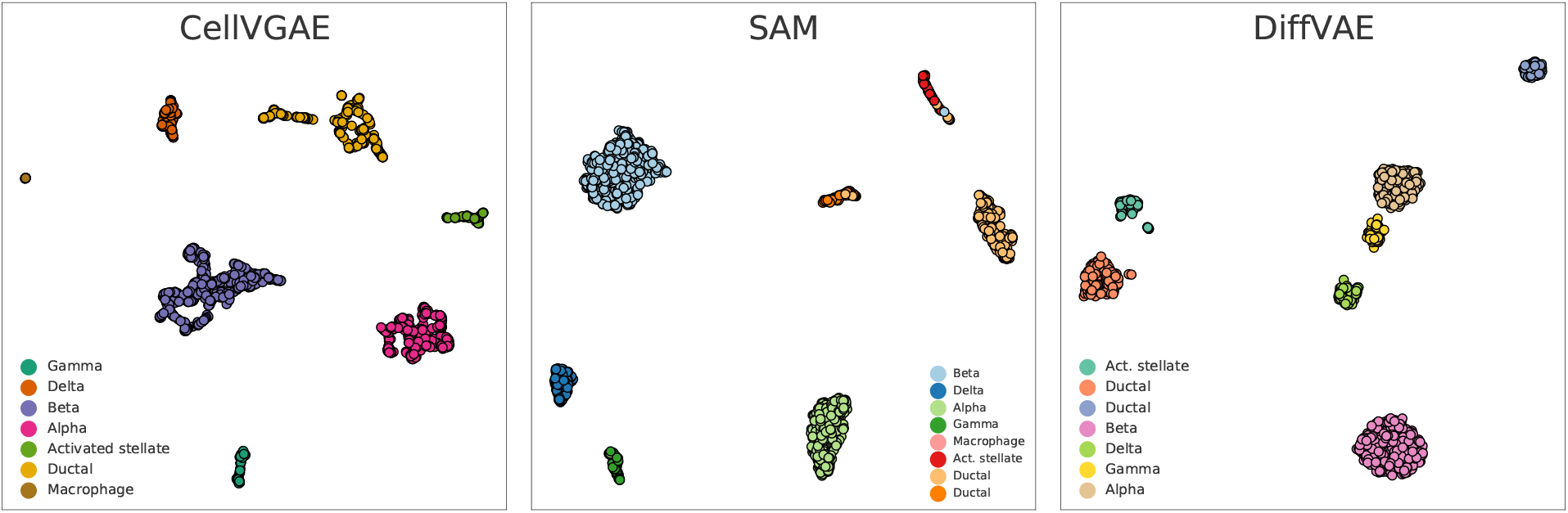
Different shape tendencies on the *Baron4* dataset.

## Footnotes

1 https://support.10xgenomics.com/single-cell-gene-expression/datasets/1.1.0/pbmc3k

2 https://satijalab.org/seurat/v3.0/pbmc3k_tutorial.html

3 https://scanpy-tutorials.readthedocs.io/en/latest/pbmc3k.html

## References

[ALB+20] Robert A. Amezquita, Aaron T. L. Lun, Etienne Becht, Vince J. Carey, Lindsay N. Carpp, Ludwig Geistlinger, Federico Marini, Kevin Rue-Albrecht, Davide Risso, Charlotte Soneson, Levi Waldron, Hervé Pagès, Mike L. Smith, Wolfgang Huber, Martin Morgan, Raphael Gottardo, and Stephanie C. Hicks. Orchestrating single-cell analysis with bioconductor. Nature Methods, 17(2):137–145, Feb 2020.

[BATCL20] Ioana Bica, Helena Andrés-Terré, Ana Cvejic, and Pietro Liò. Unsupervised generative and graph representation learning for modelling cell differentiation. Scientific Reports, 10(1):9790, Jun 2020.

[BGLL08] Vincent D Blondel, Jean-Loup Guillaume, Renaud Lambiotte, and Etienne Lefebvre. Fast unfolding of communities in large networks. Journal of Statistical Mechanics: Theory and Experiment, 2008(10):P10008, Oct 2008.

[BVW+16] Maayan Baron, Adrian Veres, Samuel L. Wolock, Aubrey L. Faust, Renaud Gaujoux, Amedeo Vetere, Jennifer Hyoje Ryu, Bridget K. Wagner, Shai S. Shen-Orr, Allon M. Klein, Douglas A. Melton, and Itai Yanai. A single-cell transcriptomic map of the human and mouse pancreas reveals inter- and intra-cell population structure. Cell Systems, 3(4):346–360.e4, Oct 2016.

[CFS09] Jie Chen, Haw-ren Fang, and Yousef Saad. Fast approximate *k*nn graph construction for high dimensional data via recursive lanczos bisection. J. Mach. Learn. Res., 10:1989–2012, December 2009.

[CNS19] Geng Chen, Baitang Ning, and Tieliu Shi. Single-cell rna-seq technologies and related computational data analysis. Frontiers in Genetics, 10:317, 2019.

[CTK+13] E. Y. Chen, C. M. Tan, Y. Kou, Q. Duan, Z. Wang, G. V. Meirelles, N. R. Clark, and A. Ma’ayan. Enrichr: interactive and collaborative HTML5 gene list enrichment analysis tool. BMC Bioinformatics, 14:128, Apr 2013.

[DCW+18] Francis Dutil, Joseph Paul Cohen, Martin Weiss, Georgy Derevyanko, and Yoshua Bengio. Towards gene expression convolutions using gene interaction graphs, 2018.

[DSL20] Jacob Deasy, Nikola Simidjievski, and Pietro Liò. Constraining variational inference with geometric jensen-shannon divergence, 2020.

[DSZ+15] Spyros Darmanis, Steven A. Sloan, Ye Zhang, Martin Enge, Christine Caneda, Lawrence M. Shuer, Melanie G. Hayden Gephart, Ben A. Barres, and Stephen R. Quake. A survey of human brain transcriptome diversity at the single cell level. Proceedings of the National Academy of Sciences, 112(23):7285–7290, 2015.

[ESM+19] Gökcen Eraslan, Lukas M. Simon, Maria Mircea, Nikola S. Mueller, and Fabian J. Theis. Single-cell rna-seq denoising using a deep count autoencoder. Nature Communications, 10(1):390, Jan 2019.

[FC16] Cong Fu and Deng Cai. Efanna: An extremely fast approximate nearest neighbor search algorithm based on knn graph, 2016.

[FL19] Matthias Fey and Jan E. Lenssen. Fast graph representation learning with PyTorch Geometric. In ICLR Workshop on Representation Learning on Graphs and Manifolds, 2019.

[GSH+15] Samantha A. Green, Mindy Smith, Rebecca B. Hasley, David Stephany, Adam Harned, Kunio Nagashima, Shahed Abdullah, Stefania Pittaluga, Tomozumi Imamichi, Jing Qin, Adam Rupert, Alex Ober, H. Clifford Lane, and Marta Catalfamo. Activated platelet-t-cell conjugates in peripheral blood of patients with hiv infection: coupling coagulation/inflammation and t cells. AIDS (London, England), 29(11):1297–1308, Jul 2015. 26002800[pmid].

[GVT+20] Christopher Heje Grønbech, Maximillian Fornitz Vording, Pascal N Timshel, Casper Kaae Sønderby, Tune H Pers, and Ole Winther. scVAE: variational auto-encoders for single-cell gene expression data. Bioinformatics, 36(16):4415–4422, 05 2020.

[HAYSZ11] Kiana Hajebi, Yasin Abbasi-Yadkori, Hossein Shahbazi, and Hong Zhang. Fast approximate nearest-neighbor search with k-nearest neighbor graph. In Proceedings of the Twenty-Second International Joint Conference on Artificial Intelligence - Volume Volume Two, IJCAI’11, page 1312–1317. AAAI Press, 2011.

[HPD19] Michael Hahsler, Matthew Piekenbrock, and Derek Doran. dbscan: Fast density-based clustering with R. Journal of Statistical Software, 91(1):1–30, 2019.

[Hun07] J. D. Hunter. Matplotlib: A 2d graphics environment. Computing in Science & Engineering, 9(3):90–95, 2007.

[KJR+16] M. V. Kuleshov, M. R. Jones, A. D. Rouillard, N. F. Fernandez, Q. Duan, Z. Wang, S. Koplev, S. L. Jenkins, K. M. Jagodnik, A. Lachmann, M. G. McDermott, C. D. Monteiro, G. W. Gundersen, and A. Ma’ayan. Enrichr: a comprehensive gene set enrichment analysis web server 2016 update. Nucleic Acids Res, 44(W1):W90–97, 07 2016.

[KW16] Thomas N. Kipf and Max Welling. Variational graph auto-encoders, 2016.

[LCK+16] Kyle M. Loh, Angela Chen, Pang Wei Koh, Tianda Z. Deng, Rahul Sinha, Jonathan M. Tsai, Amira A. Barkal, Kimberle Y. Shen, Rajan Jain, Rachel M. Morganti, Ng Shyh-Chang, Nathaniel B. Fernhoff, Benson M. George, Gerlinde Wernig, Rachel E.A. Salomon, Zhenghao Chen, Hannes Vogel, Jonathan A. Epstein, Anshul Kundaje, William S. Talbot, Philip A. Beachy, Lay Teng Ang, and Irving L. Weissman. Mapping the pairwise choices leading from pluripotency to human bone, heart, and other mesoderm cell types. Cell, 166(2):451–467, Jul 2016.

[LJKBJ17] Chieh Lin, Siddhartha Jain, Hannah Kim, and Ziv Bar-Joseph. Using neural networks for reducing the dimensions of single-cell RNA-Seq data. Nucleic Acids Research, 45(17):e156–e156, 07 2017.

[LMK20] Eugene Lin, Sudipto Mukherjee, and Sreeram Kannan. A deep adversarial variational autoencoder model for dimensionality reduction in single-cell rna sequencing analysis. BMC Bioinformatics, 21(1):64, Feb 2020.

[LMM16] Aaron T. L. Lun, Davis J. McCarthy, and John C. Marioni. A step-by-step workflow for low-level analysis of single-cell rna-seq data with bioconductor. F1000Res., 5:2122, 2016.

[LT19] Malte D Luecken and Fabian J Theis. Current best practices in single-cell rna-seq analysis: a tutorial. Molecular Systems Biology, 15(6):e8746, 2019.

[LVR15] Fong W. Lam, K. Vinod Vijayan, and Rolando E. Rumbaut. Platelets and their interactions with other immune cells. Comprehensive Physiology, 5(3):1265–1280, Jul 2015. 26140718[pmid].

[LVVD+17] Keara Lane, David Van Valen, Mialy M. DeFelice, Derek N. Macklin, Takamasa Kudo, Ariel Jaimovich, Ambrose Carr, Tobias Meyer, Dana Pe’er, Stephane C. Boutet, and Markus W. Covert. Measuring signaling and rna-seq in the same cell links gene expression to dynamic patterns of nf-kb activation. Cell Systems, 4(4):458–469.e5, Apr 2017.

[MDG+16] Mauro J. Muraro, Gitanjali Dharmadhikari, Dominic Grün, Nathalie Groen, Tim Dielen, Erik Jansen, Leon van Gurp, Marten A. Engelse, Francoise Carlotti, Eelco J.P. de Koning, and Alexander van Oudenaarden. A single-cell transcriptome atlas of the human pancreas. Cell Systems, 3(4):385–394.e3, Oct 2016.

[MDVK20] Shahin Mohammadi, Jose Davila-Velderrain, and Manolis Kellis. A multiresolution framework to characterize single-cell state landscapes. Nature Communications, 11(1):5399, Oct 2020.

[MHA17] Leland McInnes, John Healy, and Steve Astels. hdbscan: Hierarchical density based clustering. The Journal of Open Source Software, 2(11), mar 2017.

[MHM20] Leland McInnes, John Healy, and James Melville. Umap: Uniform manifold approximation and projection for dimension reduction, 2020.

[Pei14] Tiago P. Peixoto. The graph-tool python library. figshare, 2014.

[PGM+19] Adam Paszke, Sam Gross, Francisco Massa, Adam Lerer, James Bradbury, Gregory Chanan, Trevor Killeen, Zeming Lin, Natalia Gimelshein, Luca Antiga, Alban Desmaison, Andreas Kopf, Edward Yang, Zachary DeVito, Martin Raison, Alykhan Tejani, Sasank Chilamkurthy, Benoit Steiner, Lu Fang, Junjie Bai, and Soumith Chintala. Pytorch: An imperative style, high-performance deep learning library. In H. Wallach, H. Larochelle, A. Beygelzimer, F. d’Alché-Buc, E. Fox, and R. Garnett, editors, Advances in Neural Information Processing Systems 32, pages 8024–8035. Curran Associates, Inc., 2019.

[PWPW20] Christina Polasky, Franziska Wendt, Ralph Pries, and Barbara Wollenberg. Platelet induced functional alteration of cd4+ and cd8+ t cells in hnscc. International Journal of Molecular Sciences, 21(20):7507, Oct 2020.

[R C13] R Core Team. R: A Language and Environment for Statistical Computing. R Foundation for Statistical Computing, Vienna, Austria, 2013.

[RKK+19] Uku Raudvere, Liis Kolberg, Ivan Kuzmin, Tambet Arak, Priit Adler, Hedi Peterson, and Jaak Vilo. g:Profiler: a web server for functional enrichment analysis and conversions of gene lists (2019 update). Nucleic Acids Research, 47(W1):W191–W198, 05 2019.

[RSP+20] Neal Ravindra, Arijit Sehanobish, Jenna L. Pappalardo, David A. Hafler, and David van Dijk. Disease state prediction from single-cell data using graph attention networks. In Proceedings of the ACM Conference on Health, Inference, and Learning, CHIL’20, page 121–130, New York, NY, USA, 2020. Association for Computing Machinery.

[SBH+19] Tim Stuart, Andrew Butler, Paul Hoffman, Christoph Hafemeister, Efthymia Papalexi, William M Mauck III, Yuhan Hao, Marlon Stoeckius, Peter Smibert, and Rahul Satija. Comprehensive integration of single-cell data. Cell, 177:1888–1902, 2019.

[SCRE09] Jennifer M. Sowa, Scott A. Crist, Timothy L. Ratliff, and Bennett D. Elzey. Platelet influence on t- and b-cell responses. Archivum Immunologiae et Therapiae Experimentalis, 57(4):235–241, Aug 2009.

[SDG+12] Marcel H. Schulz, William E. Devanny, Anthony Gitter, Shan Zhong, Jason Ernst, and Ziv Bar-Joseph. Drem 2.0: Improved reconstruction of dynamic regulatory networks from time-series expression data. BMC Systems Biology, 6(1):104, Aug 2012.

[Sej20] Terrence J. Sejnowski. The unreasonable effectiveness of deep learning in artificial intelligence. Proceedings of the National Academy of Sciences, 2020.

[SGYP20] Valentine Svensson, Adam Gayoso, Nir Yosef, and Lior Pachter. Interpretable factor models of single-cell RNA-seq via variational autoencoders. Bioinformatics, 36(11):3418–3421, 03 2020.

[SPE+16] Åsa Segerstolpe, Athanasia Palasantza, Pernilla Eliasson, Eva-Marie Andersson, Anne-Christine Andréasson, Xiaoyan Sun, Simone Picelli, Alan Sabirsh, Maryam Clausen, Magnus K. Bjursell, David M. Smith, Maria Kasper, Carina Ämmälä, and Rickard Sandberg. Single-cell transcriptome profiling of human pancreatic islets in health and type 2 diabetes. Cell metabolism, 24(4):593–607, Oct 2016. 27667667[pmid].

[TBW+09] Fuchou Tang, Catalin Barbacioru, Yangzhou Wang, Ellen Nordman, Clarence Lee, Nanlan Xu, Xiaohui Wang, John Bodeau, Brian B. Tuch, Asim Siddiqui, Kaiqin Lao, and M. Azim Surani. mrna-seq whole-transcriptome analysis of a single cell. Nature Methods, 6(5):377–382, May 2009.

[TMSM18] Divyanshu Talwar, Aanchal Mongia, Debarka Sengupta, and Angshul Majumdar. Autoimpute: Autoencoder based imputation of single-cell rna-seq data. Scientific Reports, 8(1):16329, Nov 2018.

[TSK+20] Amirsina Torfi, Rouzbeh A. Shirvani, Yaser Keneshloo, Nader Tavaf, and Edward A. Fox. Natural language processing advancements by deep learning: A survey, 2020.

[TXL+19] Alexander J Tarashansky, Yuan Xue, Pengyang Li, Stephen R Quake, and Bo Wang. Self-assembling manifolds in single-cell rna sequencing data. Elife, 8:e48994, 2019.

[VCC+18] Petar Veličković, Guillem Cucurull, Arantxa Casanova, Adriana Romero, Pietro Liò, and Yoshua Bengio. Graph attention networks, 2018.

[VDDP18] Athanasios Voulodimos, Nikolaos Doulamis, Anastasios Doulamis, and Eftychios Protopapadakis. Deep learning for computer vision: A brief review. Computational Intelligence and Neuroscience, 2018:7068349, Feb 2018.

[vdMH08] Laurens van der Maaten and Geoffrey Hinton. Visualizing data using t-SNE. Journal of Machine Learning Research, 9:2579–2605, 2008.

[VLBM08] Pascal Vincent, Hugo Larochelle, Yoshua Bengio, and Pierre-Antoine Manzagol. Extracting and composing robust features with denoising autoencoders. In Proceedings of the 25th International Conference on Machine Learning, ICML’08, page 1096–1103, New York, NY, USA, 2008. Association for Computing Machinery.

[WAT18] F. Alexander Wolf, Philipp Angerer, and Fabian J. Theis. Scanpy: large-scale single-cell gene expression data analysis. Genome Biology, 19(1):15, Feb 2018.

[WSW+16] Yue J. Wang, Jonathan Schug, Kyoung-Jae Won, Chengyang Liu, Ali Naji, Dana Avrahami, Maria L. Golson, and Klaus H. Kaestner. Single-cell transcriptomics of the human endocrine pancreas. Diabetes, 65(10):3028–3038, Oct 2016. 27364731[pmid].

[Wtsdt20] Michael Waskom and the seaborn development team. mwaskom/seaborn, September 2020.

[WZP+17] Bo Wang, Junjie Zhu, Emma Pierson, Daniele Ramazzotti, and Serafim Batzoglou. Visualization and analysis of single-cell rna-seq data by kernel-based similarity learning. Nature Methods, 14(4):414–416, Apr 2017.

[ZSE17] Shengjia Zhao, Jiaming Song, and Stefano Ermon. Infovae: Information maximizing variational autoencoders. CoRR, abs/1706.02262, 2017.

[ZZX+20] Xiaoshu Zhu, Jie Zhang, Yunpei Xu, Jianxin Wang, Xiaoqing Peng, and Hong-Dong Li. Single-cell clustering based on shared nearest neighbor and graph partitioning. Interdisciplinary Sciences: Computational Life Sciences, 12(2):117–130, Jun 2020.

